# FYN regulates aqueous humor outflow and IOP through the phosphorylation of VE-cadherin

**DOI:** 10.1101/2023.09.04.556253

**Authors:** Krishnakumar Kizhatil, Graham Clark, Daniel Sunderland, Aakriti Bhandari, Logan Horbal, Revathi Balasubramanian, Simon John

## Abstract

The exact sites and molecules that determine resistance to aqueous humor drainage and control intraocular pressure (IOP) need further elaboration. Proposed sites include the inner wall of Schlemms’s canal and the juxtacanalicular trabecular meshwork ocular drainage tissues. The adherens junctions (AJs) of Schlemm’s canal endothelial cells (SECs) must both preserve the blood-aqueous humor (AQH) barrier and be conducive to AQH drainage. How homeostatic control of AJ permeability in SC occurs and how such control impacts IOP is unclear. We hypothesized that mechano-responsive phosphorylation of the junctional molecule VE-CADHERIN (VEC) by SRC family kinases (SFKs) regulates the permeability of SEC AJs. We tested this by clamping IOP at either 16 mmHg, 25 mmHg, or 45 mmHg in mice and then measuring AJ permeability and VEC phosphorylation. We found that with increasing IOP: 1) SEC AJ permeability increased, 2) VEC phosphorylation was increased at tyrosine-658, and 3) SFKs were activated at the AJ. Among the two SFKs known to phosphorylate VEC, FYN, but not SRC, localizes to the SC. Furthermore, FYN mutant mice had decreased phosphorylation of VEC at SEC AJs, dysregulated IOP, and reduced AQH outflow. Together, our data demonstrate that increased IOP activates FYN in the inner wall of SC, leading to increased phosphorylation of AJ VEC and, thus, decreased resistance to AQH outflow. These findings support a crucial role of mechanotransduction signaling in IOP homeostasis within SC in response to IOP. These data strongly suggest that the inner wall of SC partially contributes to outflow resistance.

## INTRODUCTION

Schlemm’s canal (SC) has a critical role in aqueous humor (AQH) drainage and intraocular pressure (IOP) homeostasis ^1–4^ SC is a flattened tube made of endothelial cells encircling the anterior portion of the eye within the tissue of the iridocorneal angle (angle formed by the iris and cornea) ^2, 5–8^. The inner wall of SC is the last barrier to AQH outflow from the eye. The ability of inner wall SECs to rapidly respond to IOP and regulate outflow is expected to be critical in achieving rapid IOP homeostasis. Abnormalities in inner wall biomechanical stiffness are likely to impact homeostatic mechanisms and IOP in glaucoma patients ^9, 10^. Despite this importance, the molecular mechanisms controlling AQH outflow through the inner wall SEC barrier are mostly undefined.

The lumen of SC is continuous to the systemic blood circulation. The inner wall of SC must preserve the blood-AQH barrier while being conductive to AQH outflow. The mechanisms underlying these incompatible functions are poorly understood ^11^. Inner wall SECs are morphologically specialized cells that express lymphatic markers like PROX1 and FLT4 ^6^. These long thin cells experience a basal to apical pressure gradient. These cells and their basement membrane provide the final barrier to AQH drainage from the eye ^1, 2, 12^. The cellular junctions in SEC are essential for maintaining barrier function.

In addition to maintaining a barrier, the cell junctions also serve as an extracellular outflow route through pores at the cell border (B-pores) ^3, 9, 13, 14^. Flow also occurs through a transcellular route where pressure-modulated intracellular pores (I-pores) in giant vacuoles form in the inner wall SECs to allow AQH outflow ^9, 15^. Supporting a role of cell junctions in determining SC permeability and outflow, blockage of intercellular clefts between SC endothelial cells (SECs) by cationic ferritin perfusion reduces outflow ^16^, while knockdown of ZO1 a tight junction associated protein increases outflow ^17^. Inhibiting PTPRB/VEPTP, a junctional receptor tyrosine phosphatase, has been recently shown to increase the outflow of AQH ^18, 19^. There is evidence that SC junctions become less complex with the overlap between cells reducing when subjected to increasing pressure ^20^. The molecular mechanisms that generally regulate AQH outflow, specifically through the cell junctions in response to IOP change, remain largely undetermined. Homeostatic mechanotransduction may control the amount of AQH outflow through intracellular and junctional pores.

Details of mechanotransduction in endothelial cells are primarily derived from studies of fluid shear stress responses of blood endothelial cells (BECs). BEC mechanotransducers that elicit flow-dependent responses include the adherens junction complex (AJC), G-protein coupled receptors, caveolae, integrins, the glycocalyx, and the cytoskeleton ^21^. In BECs, the AJC is comprised of the membrane proteins CDH5(VE-CADHERIN, VEC), PECAM1, and KDR ^22^. VEC is the primary junction molecule required to maintain barrier function by promoting homophilic adhesion in vascular endothelium, forming zipper-like AJs at cell contacts. The cytoplasmic VEC domain interacts with intracellular proteins (including regulatory protein p120-catenin,β**-**catenin that links VEC with the cytoskeleton), signaling kinases, and receptors.

Mechanical forces on cells activate SRC family kinases (SFKs). Two SFKs, SRC and FYN, are essential activators of mechanoresponses arising from the AJC complex ^22–27^. SFK phosphorylation of VEC at key tyrosine residues is required to induce junctional permeability ^28–31^. Recently, the SFK YES has been shown to regulate the plasticity of AJCs ^32^. SFK-dependent phosphorylation of VEC tyrosine 658 and 685 (pY658, pY685 VEC) in response to shear stress has been demonstrated *in vivo* in veins ^30, 31^. Phosphorylation at Y658 or Y685 induces a dynamic state of the AJ, leading to more permeable junctions in the presence of permeability-inducing agents ^30, 32^. VEC Y731 is another target of SFKs but is not required for barrier regulation in BECs^31^. The expression of mutant VECs with either Y658F or Y685F phosphorylation-resistant mutations (tyrosine substituted by phenylalanine) prevents the formation of a permeable junction in cell culture and *in vivo* ^30, 31^.

Using mice, extensive whole mount immunofluorescence imaging, and physiological methods, we show that SFK-mediated site-specific phosphorylation of VEC at SEC adherens junctions (AJs) regulates AQH outflow and IOP. SEC AJs were lined with VEC and associated cytoplasmic catenin partners. The amount of lectin flow tracer at the cell junctions (marked by VEC) increased with increasing IOP, indicating increased junctional permeability. The increased junctional permeability correlated with increased pY658VEC and activated SFK at the SEC adherens junction. FYN but not SRC is expressed in SECs, and the absence of FYN resulted in the loss of Y658 and Y685 VEC phosphorylation. Dysregulation of AQH drainage and IOP occurs in Fyn KO mice. These results indicate a role for FYN in controlling IOP and AQH outflow by controlling AJ permeability in the SEC.

## RESULTS

### SECs have continuous cell junctions with VEC and VEC-binding catenin proteins

VEC and VEC-binding proteins play a critical role in maintaining the barrier function of the AJ. These proteins also regulate AJ permeability in BECs ^29, 33^. VEC is present at the cell junctions of SC’s inner and outer walls ^6, 34^. The robust almost continuous presence of VEC marks the inner wall AJs of the SC in the eyes of a *Prox1*-GFP mouse (Top panels, Figure 1, *Prox1* is an SC marker that is most robustly expressed in the inner wall) ^6, 35, 36^. We also found that both p120-catenin and β-catenin (middle panels and bottom panels, Figure 1), which bind specific cytoplasmic domains of VEC and regulate VEC adhesivity, are co-expressed at the AJ of SECs. These results indicate that SECs have an AJ like BECs.

**Figure 1.**
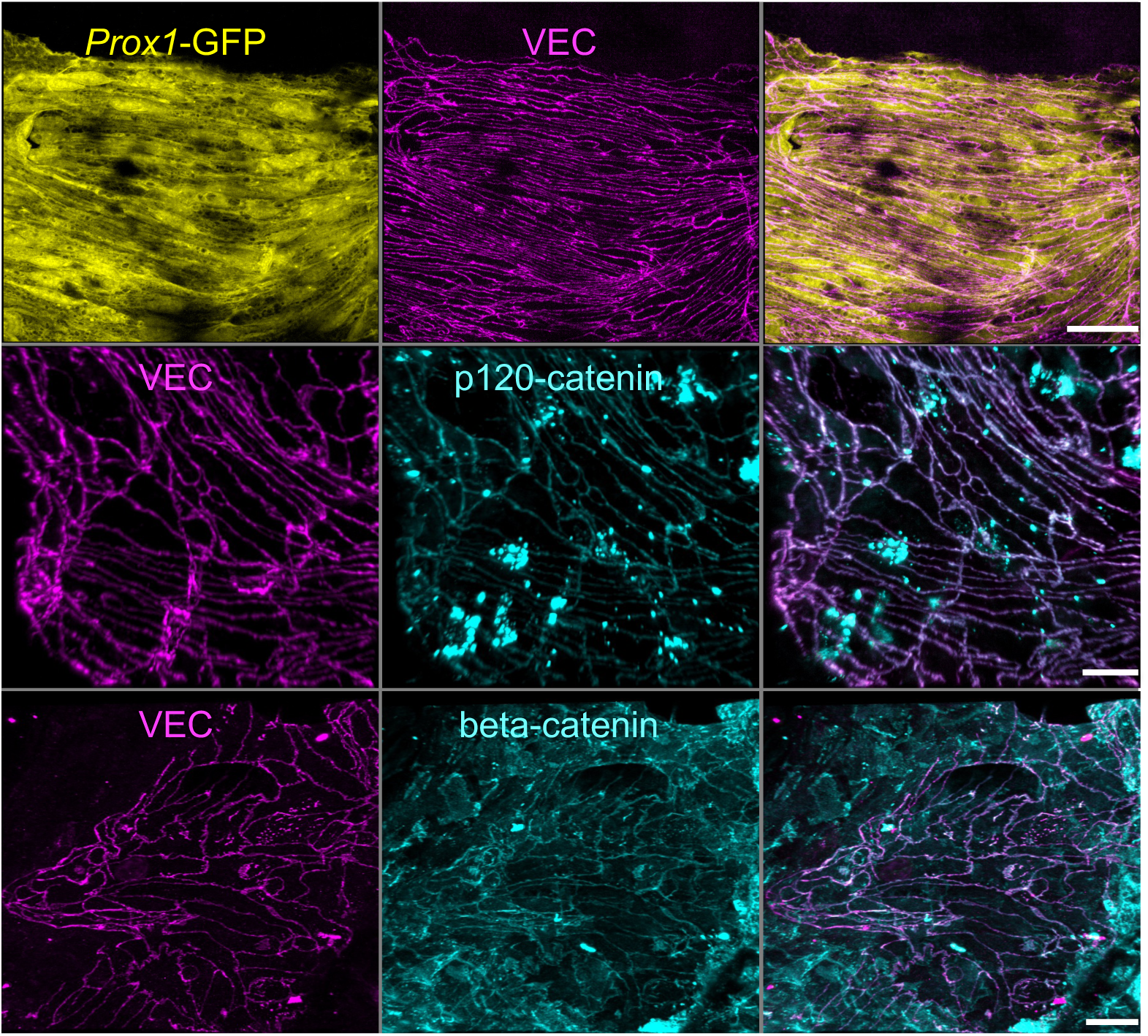
SEC AJ expresses VEC and VEC intracellular adapter proteins, p120-catenin and beta-catenin. VEC and associated cytoplasmic proteins in SC cells. Top, VEC decorates cell junctions of the SC. SC labeled with *Prox1*-GFP (yellow) and VEC (magenta). Middle, VEC labeled cell junction contains VEC associated proteins p120-catenin. Bottom, β-catenin localizes to cell junctions of SC along with VEC. N=4 eyes each. Scale bar, A-B 10µm.

### The permeability of SEC AJs increases with increasing IOP

Next, we determined if the AJs in SEC became more permeable in response to IOP elevation. We designed a protocol to perfuse *Prox1*-GFP mouse eyes in vivo with fluorescent GS lectin at set pressures (16mmHg, 25mmHg, 45mmHg) with rapid fixation of intact eyes at the set pressure using acrolein (Figure 2A). The GS lectin binds terminal, nonreducing α- and β-*N*-acetyl-D-glucosaminyl (GlcNAc) residues of glycoproteins, thus serving as stable marker for the path of AQH flow. We marked the AJ in anterior segment whole-mounts using VEC immunofluorescence. By confocal microscopy, we sampled ∼44% of the SC at high resolution (sixteen 246µm x 246µm tiles at 63x to cover 4 mm of the diameter of SC in each, n=6 eyes). The images were deconvolved, and the amount of lectin colocalizing with the junctions was calculated (Figure 2B and 2C, Figure S1). The colocalization analysis was vital because the bulk of the lectin is found in the trabecular meshwork and complicates analysis without colocalization. Compared to 16mmHg IOP, we observed a 3.6-fold and 5-fold increase in lectin fluorescence at the AJ at 25 and 45 mmHg, respectively. Thus, AJ permeability in the inner wall of SC increases with increasing IOP.

**Figure 2.**
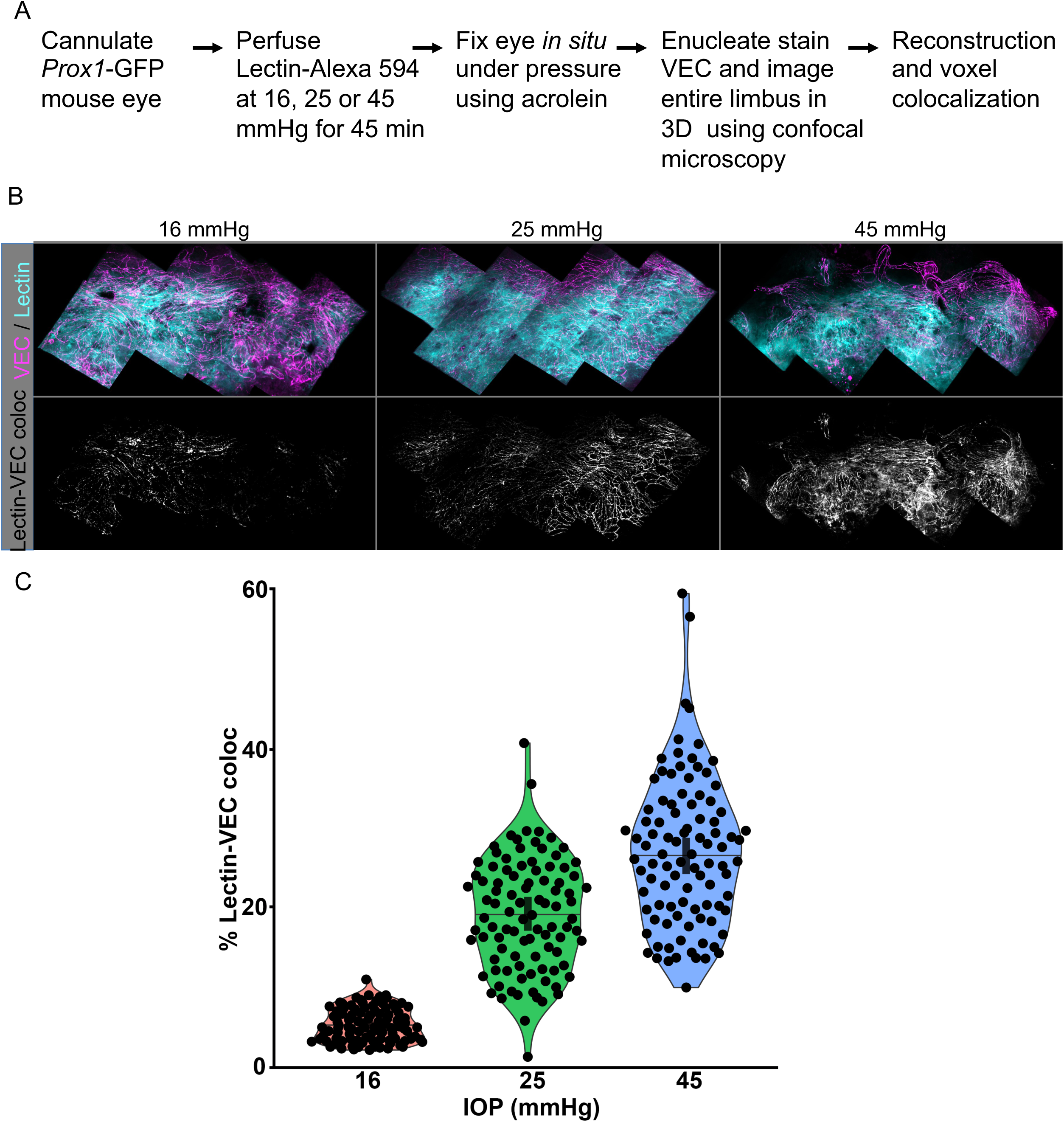
Lectin tracer levels increase at cell junctions with increasing IOP. **A.** Experimental scheme, **B.** *Prox1*-GFP eye perfused with lectin at 16, 25 or 45 mmHg. Top panels, SC’s inner wall showing VEC and lectin labeling, Bottom panels, colocalization of VEC and lectin at the AJ. Fig S1 shows all proteins imaged separately. **C.** Quantification of lectin-VEC colocalization within the inner wall of SC in eyes perfused at 16 mmHg (mean ± SD, 5.3±2.1 %), 25 mmHg (mean ± SD, 19.1±7%) and 45 mmHg (mean ± SD, 26.9± 9.3%). Each of the 96 points represents data for the entire width of a 246µm lengths of SC from each quadrant (4 tiles per quadrant x 4 quadrants x 6 eyes) at each pressure. One-way ANOVA results, F (2,285) = 244.3, p<0.00001. Horizontal line in violin plot is the median and vertical bar the 95%CI. Scale, 25µm.

### Destabilizing phosphorylation of VEC Y658 residue with increasing IOP

Phosphorylation of VEC at key tyrosine residues destabilizes junctional interactions and is required for induction of junctional permeability ^28–31^. Phosphorylation of VEC tyrosine, pY658 and pY685, in response to shear stress occurs *in vivo* in veins ^30, 31^. Given that SC originates from veins ^6, 37^, it is likely that the AJ permeability in SEC results from phosphorylation of Y658 and Y685 in the VEC cytoplasmic domain.

To determine if IOP elevation resulted in phosphorylation of pY658 and pY685, we clamped IOP in living mice at 16 mmHg, 25 mmHg, or 45 mmHg for 45 min and then rapidly fixed the eyes *in situ* under pressure (Figure. 3A). We determined the extent of VEC tyrosine phosphorylation at Y658 and Y685 in the SC using phospho-specific antibodies on whole-mounts ^30^. We analyzed the amount of phospho-VEC colocalized with the AJ labeled using a VEC antibody and sampling about 44% of SC (as above). Compared to 16 mmHg, AJ VEC phosphorylation increased 1.4-fold at Y658 (indicated as pY658) at 25mmHg and 3.6-fold at 45mmHg (Figure. 3B and C, Figure S2). In contrast, in the same time frame, pY685 was detected at the AJ at 16 mmHg but was significantly decreased at higher IOP, suggesting dephosphorylation of pY685 at 25 mmHg and 45 mmHg (Figure 4). These results show that IOP elevation results in the phospho-regulation of AJ resident VEC at Y658 and Y685.

**Figure 3.**
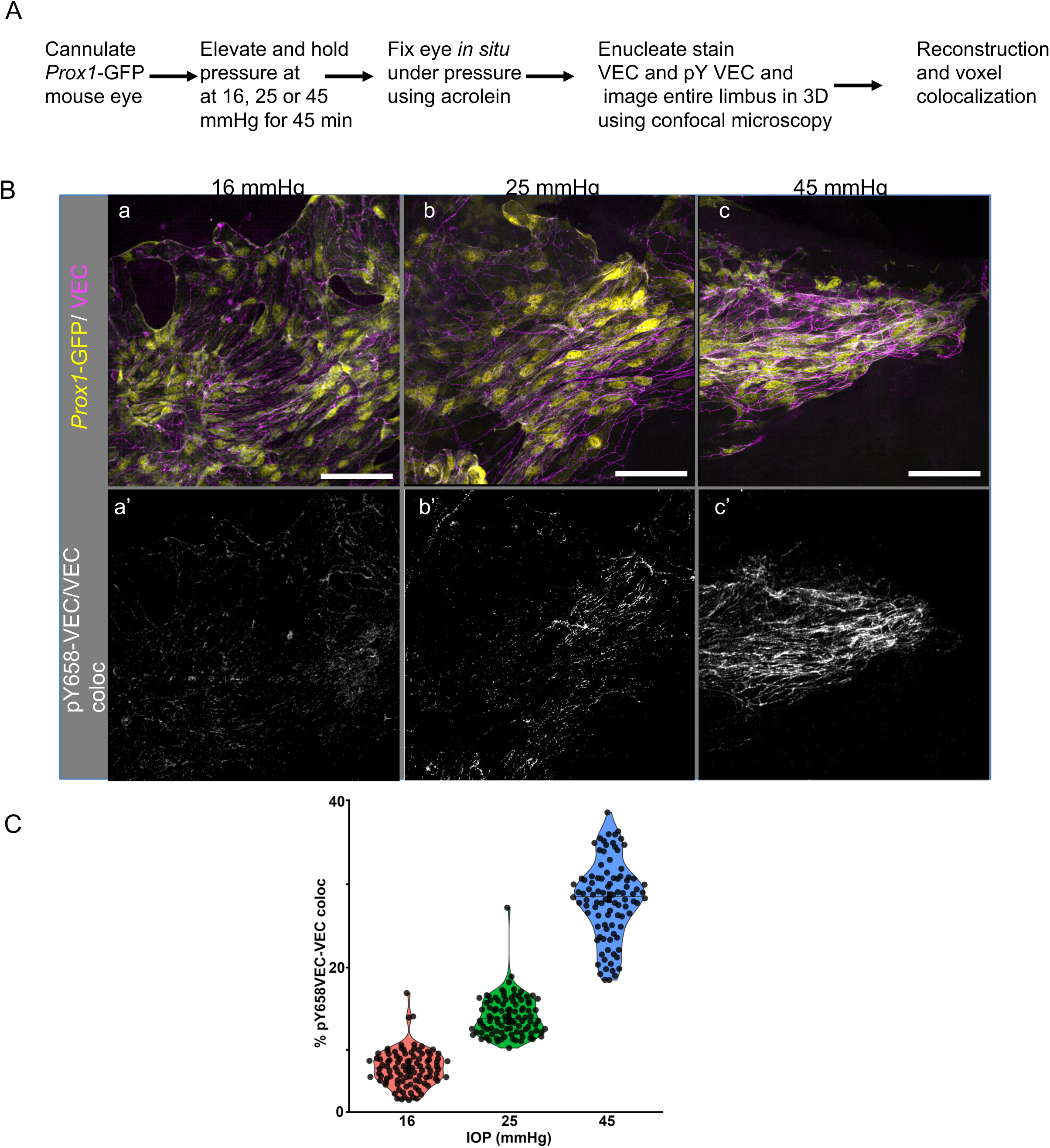
Site-specific tyrosine phosphorylation (pY658) of VEC increases with increasing IOP. **A.** Experimental scheme, **B.** *Prox1*-GFP eye pressure clamped at 16 mmHg, 25 mmHg, or 45 mmHg. **a-c**, the inner wall of SC with GFP and VEC labeling, and **a’-c’** colocalization of pY658VEC and VEC at cell-cell junctions at the indicated pressure. C. Quantification of pYVEC-VEC colocalization within the inner wall of SC in eyes perfused at 16 mmHg (mean ± SD, 7.8±2.2 %), 25 mmHg (mean ± SD, 14±2.4%) and 45 mmHg (mean ± SD, 28±4.6%). Each of the 96 points represents data from a 246µm length of SC at each pressure (4 tiles per quadrant x 4 quadrants x 6 eyes. One-way ANOVA results, F (2,285) = 947.1, p<0.00001. Horizontal line in violin plot is the median and vertical bar the 95%CI. FigS2 shows all proteins visualized as individual channels. Scale, 20µm

**Figure 4.**
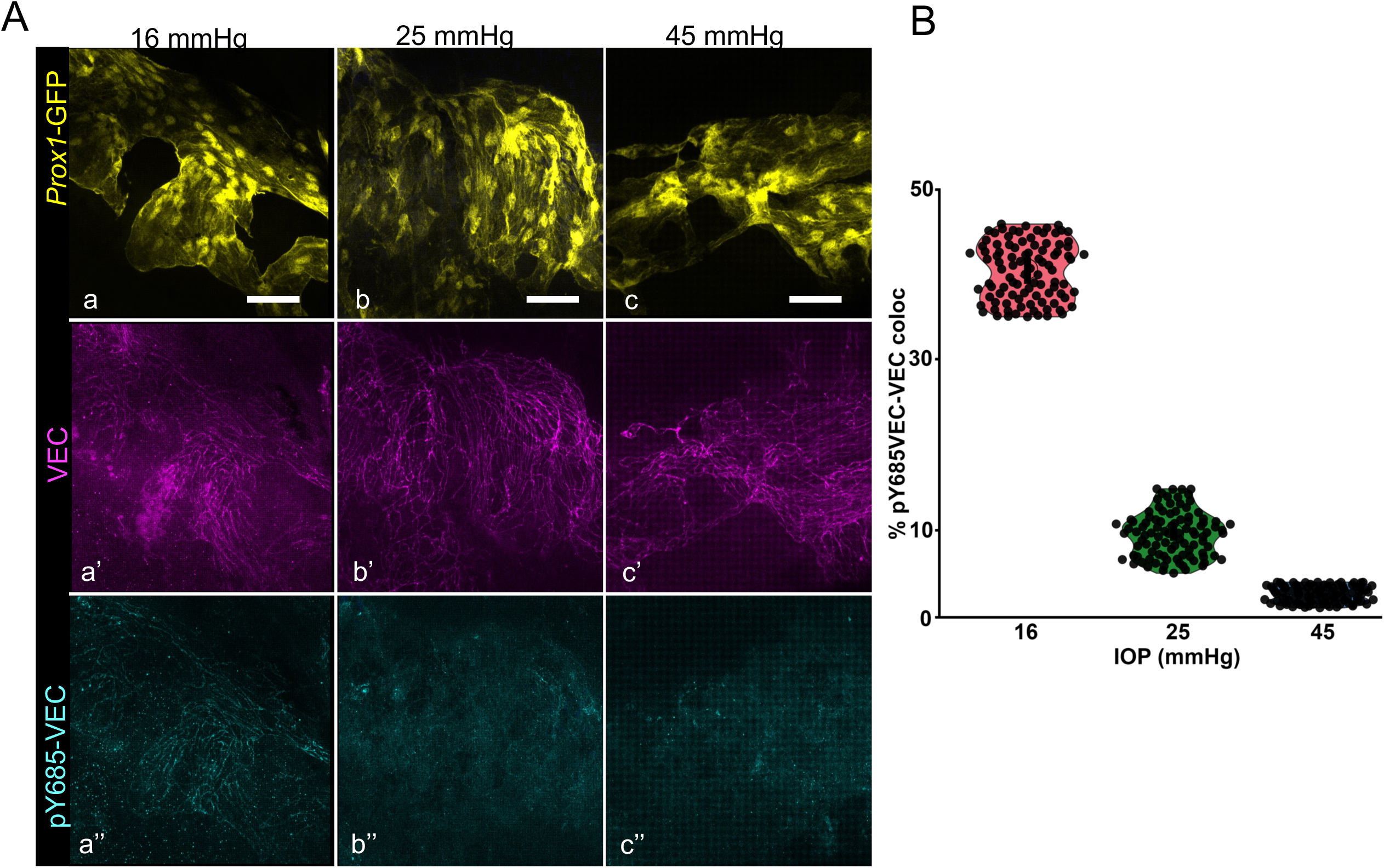
VEC Y685 tyrosine phosphorylation decreases with increasing IOP. **A**. *Prox1*-GFP eye pressure clamped at 16 mmHg, 25 mmHg, and 45 mmHg. **a-c,** inner wall of SC showing GFP and VEC labeling, **a’-c’** VEC and **a’’-c’’** localization of pY658VEC at cell-cell junctions at the indicated pressure. **B**. Quantification of pYVEC-VEC colocalization within the inner wall of SC in eyes perfused at 16 mmHg (mean ± SD.= 40.4±3.4 %), 25 mmHg (mean ± SD.= 9.6±2.6%) and 45 mmHg (mean ± SD= 2.6±0.92%). Each of the 96 points represents data from a 246µm length of SC from each quadrant (4 tiles per quadrant x 4 quadrants x 6 eyes) at each pressure. One-way ANOVA results, F (2,285) = 947.1, p<0.00001. Scale, 20µm.

### Increasing IOP activates SFKs at SEC inner wall AJs

SFKs phosphorylate tyrosine residues in the VEC cytoplasmic domain. SFK-dependent phosphorylation of VEC Y658 and Y685) is demonstrated in response to shear stress in vivo in veins ^30, 31^. Therefore, SFKs are strong candidates to phosphorylate VEC at AJs in the inner wall of Schlemm’s canal. SFKs activation occurs upon phosphorylation at SFK Y418 ^38, 39^. To assess SFK activation at cell junctions in the inner wall of SC, we determined the levels of activated (phosphorylated) SFK at different pressures by immunostaining with a pan SFK, pY418 specific antibody ^40, 41^ (Figure 5). We elevated IOP in mouse eyes and rapidly fixed eyes under pressure, prepared whole-mounts, and immunostained for pY418SFK and VEC (Figure 5A). With increasing IOP, the pY418 SFK signal increased at SC inner wall cell junctions (Figure 5B and C). Compared to 16 mmHg, SFK activation increased 2.9-fold at 25 mmHg and 3.7-fold at 45 mmHg. Thus, an increase in IOP results in an increase in SFK activation.

**Figure 5.**
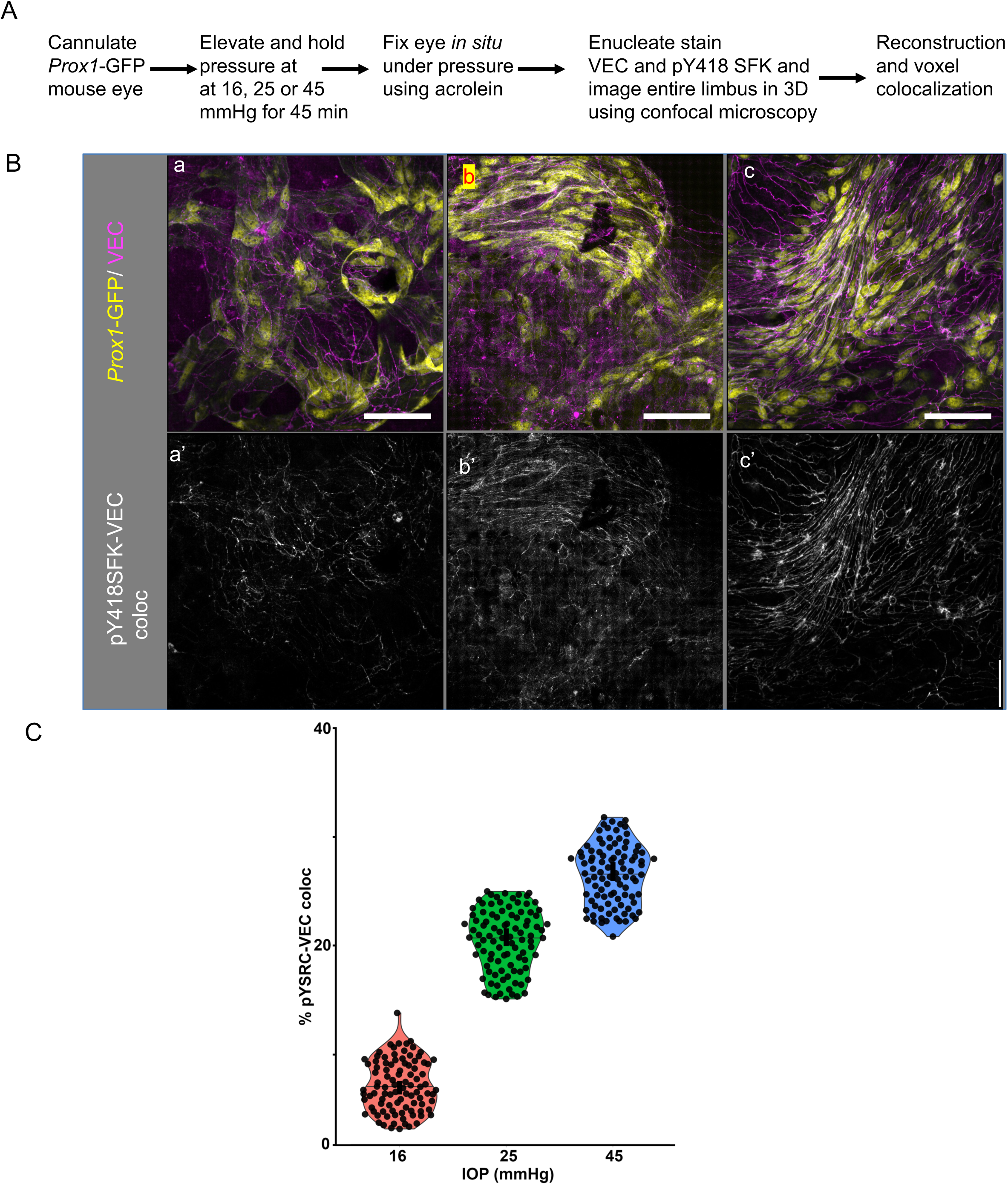
SRC family kinases are activated by increasing IOP. A. Experimental scheme, B. *Prox1*-GFP eye pressure clamped as indicated (need show pressures above panels). a-c, the inner wall of SC showing GFP and VECAD labeling, and a’-c’ colocalization of pY418SFK and VECAD at cell-cell junctions. C. Quantification of pYVECAD-VECAD colocalization within the inner wall of SC in perfused eyes (Pressure, mean ± SD; 16 mmHg,7±2.3%, 25 mmHg, 20.4±2.9%; and 45 mmHg, 26.5±2.8%. Each of the 96 points as in previous figures. One way ANOVA results, F (2,285) = 1346.2, p<0.00001. Horizontal line in violin plot is the median and vertical bar the 95%CI. Scale, 20µm.

### FYN is expressed in SECs

Two SFK’s, SRC and FYN, are established to phosphorylate tyrosine residues in the cytoplasmic domain of VEC ^28, 30, 42, 43^. By analyzing our recently generated single-cell sequencing data for limbal tissue, we show that *Fyn* is expressed in SECs (including inner wall SECs) while *Src* is not and confirmed this by IHC of anterior segment whole mounts (Figure 6, Figure S3, S4) ^44^. These data are similar to those provided in a recent, single-cell sequencing analysis ^45^. Since FYN is the SFK expressed in SC, we continued to determine the requirement of FYN for normal IOP regulation.

**Figure 6.**
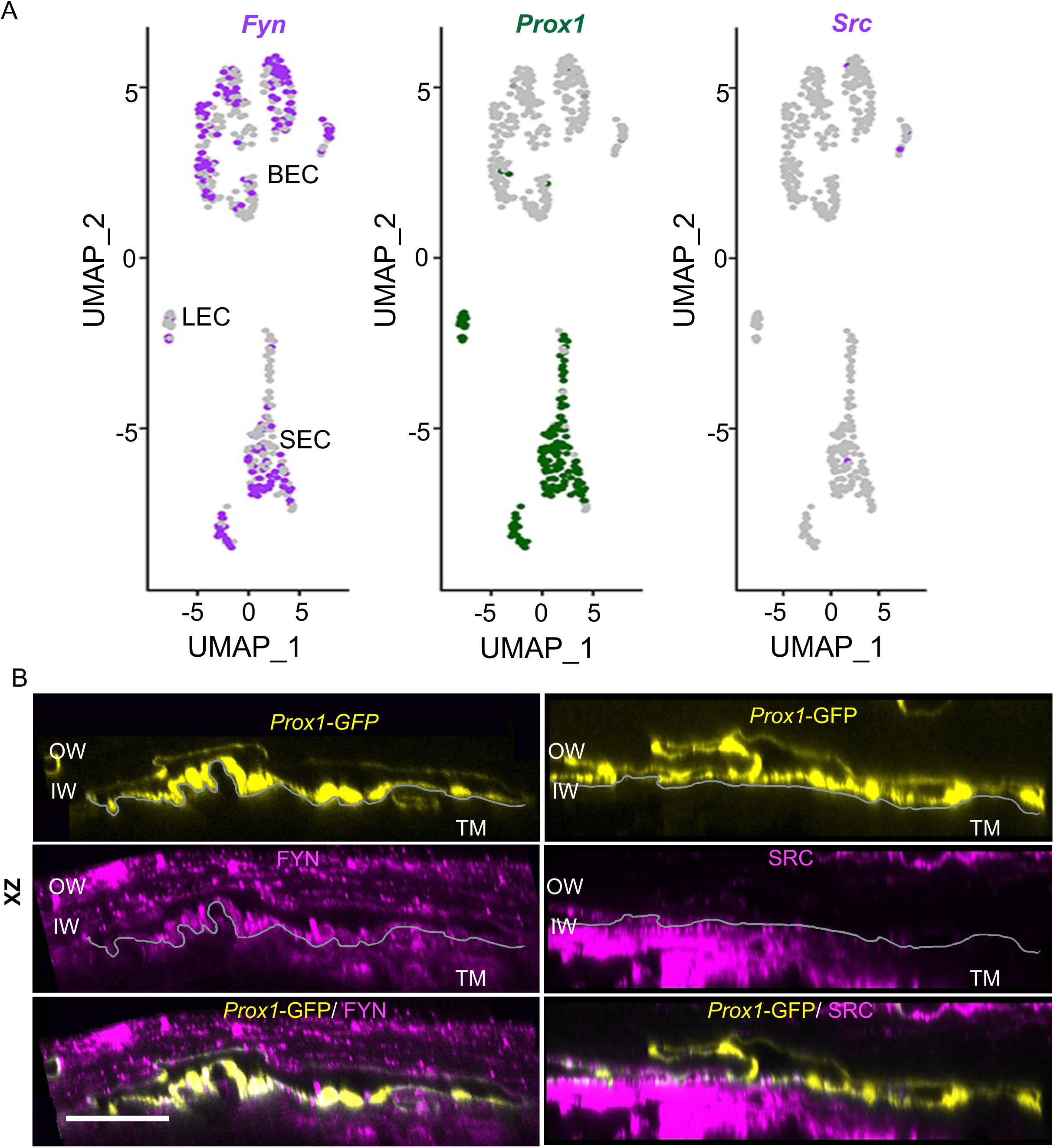
SFKs capable of phosphorylating VEC are present in SC and TM. **A.** Feature plot representation of limbal endothelial cells obtained from single cell RNA sequencing shows *Fyn* but not *Src* is expressed in the *Prox1* expressing SECs. *Fyn* also localizes to blood endothelial cells in contrast to *Src*. **B**. XZ confocal sections of *Prox1*-GFP (yellow) eyes encompassing SC and TM show that FYN localizes (magenta, left) predominantly to the SC while SRC (magenta, right) localizes predominantly to the TM. The gray line marks the border between SC IW and TM in first and second. XZ optical sections were sliced along the section of XY show in Fig S4. N=4 eyes. Scale, 20µm.

### FYN phosphorylates VEC in the inner wall of SC

To find the functional role of FYN in the SC we determined the effects of loss of FYN at a molecular and physiological level. We started by immunostaining with a VEC antibody. The general morphology of SC in FYN mutant (*Fyn* ^-/-^) mice was indistinguishable from SC in control (WT) mice (Figure 7A). At a light microscopy level, the iridocorneal angle of the WT and *Fyn* ^-/-^ appear similar (Figure S5). At an ultrastructure level, electron microscopy showed that the SC cell junctions and TM looked similar in WT and FYN mice (Figure S6). Next, we determined if the loss of FYN resulted in changes to the phosphorylation pattern of VEC. We performed immunoblotting on lysates obtained from limbal strips of wild type and FYN null eyes (Figure 7B and Figure S5). FYN, as predicted, was absent in FYN null mice but present in control at a band close to the expected size of 59-kDa (Figure 7B and Figure S7). VEC expression is at similar levels in control and FYN null mice (Figure 7B and Figure S7). pYVEC658 band was <90% of the control band after normalization of band intensity to GAPDH levels. pY685 VEC was almost wholly absent in mutants. The basal level of SFK activation indicator pY418 SRC was also reduced by ∼72 % in the FYN null lysate. Immunofluorescence showed the loss of pY658 and pY685 VEC at AJ in SEC. Activation of SFK, as judged by pY418SRC phosphorylation, was reduced in the Fyn mutant SC. These results show that FYN is the primary SFK responsible for phosphorylating VEC in the inner wall of SC.

**Figure 7.**
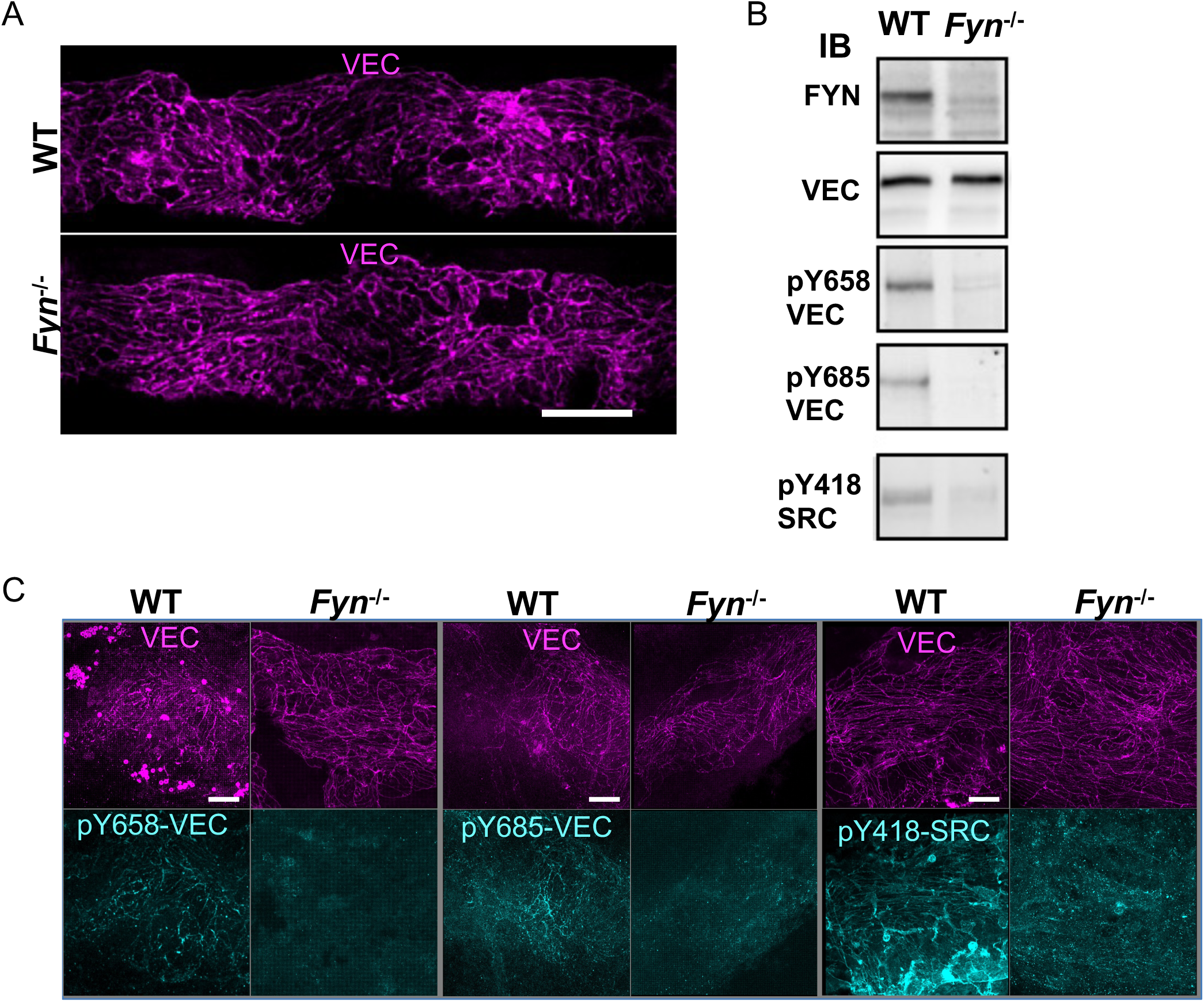
Loss of FYN results in loss of VEC phosphorylation in SEC. A. Loss FYN does not affect the gross SC morphology, N=6 eyes B. Comparison of WT and *Fyn^-/-^* limbal strip lysates (pool of 6 eyes) by immunoblotting shows the reduction pf pY658VEC and pY418 SRC levels with loss of FYN. Antibodies used for immunoblotting are shown to the left of the blots under the heading IB. pYVEC658 in control was represented by a single band around the expected 120-KDa size. A faint doublet band was present in the FYN null that was slightly larger and slightly smaller than the pY658 VEC band in WT. C. FYN null SC shows loss of phosphorylated VEC and activated SRC. pY658 VEC (left), pY685VEC (middle), and pY418 SRC (right), Phosphoproteins are shown in cyan and VEC in magenta. N=3 eyes for each genotype and antibody. Scale, A, 50 µm and E, Scale, 20µm.

### FYN null mice have dysregulated AQH outflow and IOP

To determine the functional role of FYN in AQH drainage and IOP regulation, we measured IOP and determined outflow facility (AQH flow out of the eye nl/min/mmHg), the standard measure of resistance to AQH drainage (Figure 8). FYN mutants have a dysregulation of IOP with an increased spread of values in each direction. Given this spread, we calculated the variance (VAR) and mean absolute deviation (MAD, see methods) and found that IOP in FYN null eyes showed increased variance (VAR=7, MAD=1.86) with a range that was 4.4-fold greater than that of controls (VAR=1.6, MAD=1.06) (Figure 8A). We saw a significant elevation of IOP in FYN mutants when comparing the third quartiles of each genotype (p=2.2x10^-7^) while IOP was significantly lower for the first quartile (p=1.5x10^-7^) and same in second quartile (p=0.85) compared to wild type mice. The dysregulation of IOP with changes in each direction resembles the spread in IOP values often detected in mice with mutations in various genes that increase IOP and induce glaucoma ^46–48^. This spread is not well understood but typically starts at ages when high IOP first becomes elevated and likely reflects abnormalities in diurnal and homeostatic control that result from the IOP elevating effects of increased resistance to aqueous humor outflow ^48^. IOP changes did not correlate with age (Figure S8A). These results show that the FYN mutation resulted in a profound dysregulation of IOP consistent with increased resistance to aqueous humor outflow.

**Figure 8.**
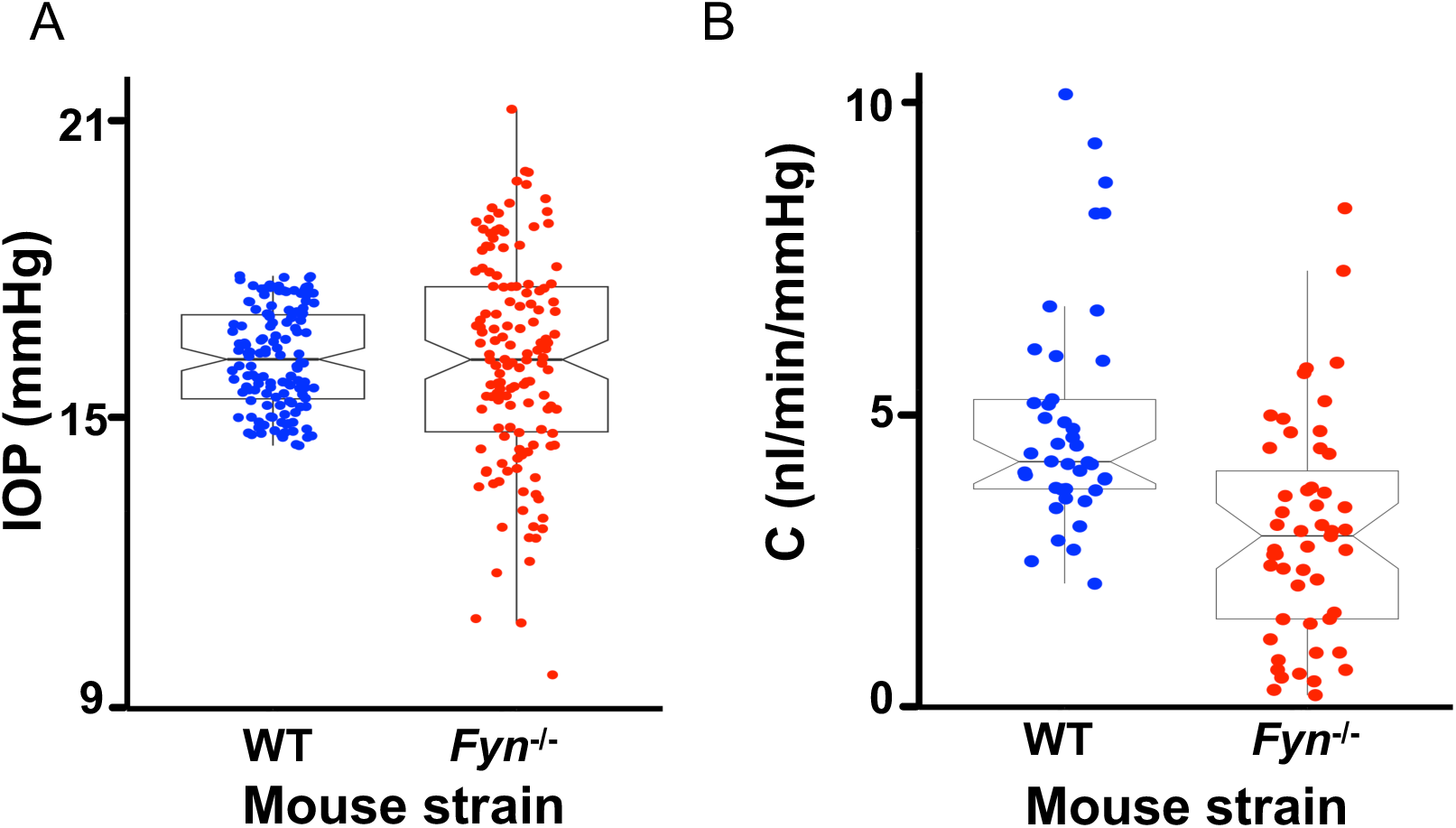
FYN is required for control of AQH outflow and IOP. **A.** IOP is dysregulated in eyes of *Fyn^-/-^* mice. The variance of IOP data for FYN null eyes was 4.4-fold that of control (7 vs 1.6) reflecting the dysregulation of IOP, N=138 eyes. 35 female and 34 male mice of 4-20 months of age. **B.** *Fyn^-/-^* mice have reduced aqueous humor outflow facility, C. *In vivo* outflow facility measurements: WT= 4.44 ±1.7and *Fyn^-/-^* ^=^ 2.75 ± 1.7, p=2.2x10^-6^, (mean ± SD) N=52 eyes, 13 female and 13 male mice of 4-20 months of age.

Unlike IOP, the outflow facility is a direct measure of the effect of the mutation on the resistance to AQH drainage at the level of TM/SC. Its measurement is subject to fewer homeostatic mechanisms than IOP as it is not dependent on the rate of AQH humor production and is free from factors that act to regulate IOP at sites away from the drainage structures. AQH outflow facility was significantly lower in FYN null eyes compared to wild type overall (Figure 8B, WT= 4.4 ±1.7 and *Fyn^-/-^* = 2.75 ± 1.7 nl/min/mmHg, p= 2.2x10^-6^) and in each of the quartiles (Q1, WT= 2.96±0.5 vs. *Fyn^-/-^* 0.8±0.4 nl/min/mmHg, p = 4.3 x 10^-13^, Q2, 4 ± 0.4 vs. 2.7 ± 0.5 nl/min/mmHg, p = 1.1 x 10^-13^ and Q3, 6.7 ± 1.6 vs. 5.0 ± 1.2 nl/min/mmHg, p=0. mean ± SD and p-value shown). Outflow changes did not correlate with age (Figure S8B). These data demonstrate that FYN is a crucial regulator determining the resistance to aqueous humor drainage.

In summary, our data show that the FYN SFK regulates IOP and AQH outflow by impacting the equilibrium between tyrosine phosphorylation and dephosphorylation of VEC at AJ in the inner wall of SC. Normal regulation of IOP requires FYN function.

## DISCUSSION

This study shows the requirement for the SFK FYN in IOP regulation. FYN responds to IOP elevation by phosphorylating specific residues in VEC’s cytoplasmic tail, which regulates SC permeability to AQH. Our data demonstrate that FYN is a crucial regulator determining the resistance to AQH drainage within the inner wall of SC. This inner wall function is pertinent as the mechanisms controlling this resistance are not clearly defined and because of the debate regarding where the key site(S) of resistance lie: within either the trabecular meshwork, SC, or possibly distal blood vessels to which SC connects ^3, 49, 50^. Our data clearly implicate the inner wall of SC as at least partially contributing to this resistance. These findings are important for understanding IOP regulation and have direct clinical relevance for developing IOP-lowering treatments to combat glaucoma, a widespread neurodegenerative disease for which IOP lowering is the standard of care.

The SC inner wall must function as a blood-AQH barrier and be permeable to AQH outflow in response to pressure. The AJ is vital in maintaining barrier function in veins ^51^, and SC is venous-derived ^6^. SC cells must sense IOP changes and, in a highly regulated manner, become permeable to AQH outflow so as not to lose their barrier function. How this is accomplished is still largely unclear. In addition to a physical barrier, the AJ complex (consisting of VEC, PECAM1, and KDR) also functions as a mechanosensor in BECs ^22^. SFKs are critical in this mechanosensing function ^22^. Thus, AJs in SC could function as sensors of pressure-dependent mechano-stimuli and responders that rapidly modulate local permeability. Such a self-contained system would allow for rapid, dynamic, local responses without compromising barrier function. AJ of SC, thus, is a strong candidate for regulating outflow.

Our study is the first to show a functional role for the AJ in SC, particularly the essential AJ adhesion protein (VEC) in regulating IOP. VEC is required for blood vessels and lymphatics to develop correctly ^52–54^ and is central in controlling vascular permeability in these tissues ^53, 54^. Our data add SC to this list of vessels, but further experiments are needed to fully understand the roles of VEC in SC and its total contribution to outflow resistance (including the specific deletion of VEC in SECs) ^54, 55^.

For SECs, phosphorylation at Y658 and Y685 was present at “normal IOP” (16 mmHg) with a marked increase of SFK activation and Y658 phosphorylation with increasing IOP. In contrast, phosphorylation at Y685 was detected only at 16 mmHg. This finding agrees with the observation that pY658 and pY685 VEC are present at basal levels in the veins and showed altered levels in response to permeability-inducing substances like bradykinin ^30, 56^. However, our finding contrasts another study showing increased Y685 phosphorylation on increasing vascular permeability through VEGF or histamine treatment (with pronounced vascular leakage) ^31^. These differences may be due to differences in a cellular context or other experimental conditions ^30, 31^The kinetics of phosphorylation of Y658 and Y685 seems to vary with the biological system studied and also the stimulus that induces phosphorylation ^30, 31, 57^. In SEC, it is also possible that phosphorylation of one of the tyrosines (e.g., pY658)^58^) reflects the mechanosensor function of AJ and the phosphorylation of the other (pY685) mediates the response to the IOP mechano-stimulus i.e., to increase junctional permeability Future studies are needed to further define the role of site-specific VEC phosphorylation(s) in regulating SC permeability and aqueous humor outflow, including the generation and use of mice with point mutations that abrogate phosphorylation at key residues ^31^.

Alterations of junctional permeability following phosphorylation of VEC at Y658, and Y685 in SEC could result from 1) The loss of p120-catenin binding. Phosphorylation of pY658, which is within the p120-catenin binding site, reduces the binding of p120-catenin and adhesivity of VEC junctions ^58–60^. However, this loss of binding of p120-catenin to pY658 VEC has not been demonstrated in BEC *in vivo* in mice or by us in this study. To correlate an association or lack therefore of p120 catenin with pY658 VEC in SEC with degrees of AJ permeability, real-time determination of p120-catenin binding to VEC along with the use a tracer to monitor changes in permeability will be needed. 2) The endocytosis of the phosphorylated VEC. We could not see a dramatic increase in levels of cytoplasmic VEC signal in most cells with the antibodies that we used. However, if sub-micron scale regions showed rapid endocytosis and recycling, we would not have detected the event. These possible mechanisms are currently experimentally inaccessible due to the limitations of technologies. In addition, compared to BEC, SEC could have a different phosphorylation threshold at which the junctions become permeable due to their unique molecular phenotype ^6^.

Regardless of the mechanism of permeability induction at the SEC AJ, an essential requirement is that SC’s barrier function is not compromised. One way to accomplish this is rapidly turning off phosphorylation events by a cognate phosphatase. pY658 and pY685 in VEC are targets of PTPRB/VE-PTP ^30, 31^. PTPRB is active in SEC ^18, 61^). The loss of pY685 could result from PTPRB activation. In a real-world scenario, this is an attractive mechanism as the phosphorylation of VEC serves as a rapid on-off switch to control AJ permeability preventing catastrophic loss of barrier function in response to rapid changes in IOP.

SFKs SRC, FYN, and YES are important activators of mechanoresponses arising from the AJC complex ^22–26, 32^. Mechanical forces on cells such as shear stress and strain can activate SFKs ^22, 62, 63^. AJ mechanoresponses to shear stress can activate SFK downstream of the mechanosensor PECAM1 ^22^. The biomechanics of the AQH drainage pathway offers some insight when considering our data. SC lumen narrows upon IOP elevation, increasing the shear stress on SECs. In contrast, TM expands with a concomitant reduction of shear stress on TM cells ^64–66^. Thus, SFK activation in SC could result from shear stress. SFKs can be activated rapidly (<0.3s) by the mechanical strain as occurs in cultured cells subjected to rapid deformation of the actin and microtubule cytoskeleton ^24, 67^. The deformation of the cytoskeleton of the long thin SEC when subjected to mechanical strain, resulting from IOP elevation, could thus result in an SFK activating event. Other possible mechanisms of SFK activation are: 1) Activation of integrins as a result of IOP elevation-related mechanical stimulus ^68, 69^, 2) Disruption of caveolae through mechanical stretching of the cell membrane, which can cause the release of SFKs from caveolin resulting in activation of SFKs ^67, 70–72^. Caveolin modulates IOP, and SNPs within C*AV1* are associated with human glaucoma ^73–75^. 3) SFKs can be activated by KDR ^29^, which in turn can be activated in SECs by VEGF secreted in response to the mechanical strain experienced by trabecular meshwork (TM) cells ^76, 77^. IOP elevation thus could activate SFKs in multiple ways.

One limitation of our study is that the FYN mutant mice had a body-wide knockout. Although we have shown the regulation of SC AJs by FYN and can explain the effects of the FYN mutation on IOP and outflow physiology, we cannot rule out at least a partial role for FYN in the TM as it is expressed) in some TM cells (albeit at a lower level). The role of SFK in the TM and its contribution to IOP regulation remains to be determined. Further measurements of IOP and AQH facility after SEC-specific knockout of FYN is needed to confirm a role solely in SECs. In addition, more work needs to be done to understand integration of other mechanisms, such as the NOS3/ NO and TIE2/ANGPT, with the FYN mediated pathway in regulating AQH outflow.

In conclusion, we have found an IOP stimulus-induced activation of FYN that regulates the permeability of the AJ by phosphorylation of VEC. These results suggest potential targets for therapy of elevated IOP, including the SFK and phosphatases (e.g., PTPRB) that regulate the phosphorylation states of VEC at the SEC AJ.

## ACKNOWLEDGEMENTS

We are grateful to Dr. Elisabetta Dejana (Uppsala University) and Fabrizio Orsenigo (FIRC Institute of Molecular Oncology: Milan) for their generous gift of the phosphor-VE-CADHERIN antibodies. Dr. Dan Stamer (Duke University) for critically reading the manuscript. The project was funded by National Eye Institute (NEI) grant R01EY032062 (KK and SWMJ), with partial contributions from R01EY028175 (KK), R01EY032507 and R01EY018606 (SWMJ). The content is solely the responsibility of the authors and does not necessarily represent the official views of the National Institutes of Health. Additional support was provided by Bright Focus Foundation grant BFOCUS CG2020004 (KK and SWMJ) and startup funds at Columbia University, including the Precision Medicine Initiative. Initial funding was provided by HHMI (SWMJ). The project was also supported by a Vison Core grant P30 EY019007 and an unrestricted departmental award from RPB.

## MATERIALS AND METHODS

### Mouse Strain, Breeding, and Husbandry

This study was performed per the guidelines in the Guide for the Care and Use of Laboratory Animals of the National Institutes of Health. All the animals were handled according to approved institutional animal care and use committee (IACUC) protocols (#99108, #21016) of The Jackson Laboratory or Columbia University (AABE9554). In addition, all experiments were conducted in accordance with the Association for Research in Vision and Ophthalmology’s statement on the use of animals in ophthalmic research. All mice were housed in a 14-h light to 10-h dark cycle under previously described conditions ^78^. Institutional Animal Care and Use Committee approved all procedures described here. After backcrossing to C57BL/6J (B6) 10 generation, the following transgenic and knockout mice were used: B6.*Prox1-GFP I,* B6*. Fyn ^tm1^ ^Sor^/*J (from Jackson Laboratory Stock no. 012468) ^79^, B6*. Src ^tm1^ ^Sor^/*J (from Jackson Laboratory Stock no. 002277) ^80^.

### Immunofluorescence on Anterior Eye-Cups

Immunofluorescence on anterior eye segments was done as described in an earlier study ^6^. Briefly, anterior eye-cups were incubated with 3% bovine serum albumin and 1% Triton X-100 in 1×phosphate buffered saline (PBS, blocking buffer) at room temperature for one h in 2 ml glass vials to block nonspecific binding of antibody and to permeabilize the tissue. The anterior cups were then incubated with primary antibodies of choice in 200 µl blocking buffer for 2 d, with rocking, at 4 °C. The anterior cups were washed three times over a 3-h period with 1× PBS containing 0.1% Tween-20. The primary antibodies were detected with the appropriate species-specific secondary antibody (all Alexa 488, 594, or 647 at 1:1,000 dilution, Thermo Fisher Sci, Waltham, MA) diluted in blocking buffer also had DAPI to label nuclei. The immunostained eyecups were washed four times over a 3-h period in 1× PBS. Eye cups were then whole mounted on slides in Prolong Diamond (Thermo Fisher Sci, Waltham, MA). The following primary antibodies were used in this study: Goat IgG against VE-cadherin (1:100, AF1002, R&D Systems, Minneapolis, MN), pY658 and pY685 VEC (1:40, gift of F. Orsenoigo and E.Dejana), pY658 VEC (1:50, 44-1144G, Thermo Fisher Sci, Waltham, MA), SRC (1:50, ab109381, Abcam, Cambridge, MA), FYN (1:50, ab125016, Abcam, Cambridge, MA), FYN (1:50, HPA023887,SIGMA-ALDRICH, Burlington, MA), pY418SRC (1:50, ab4816, Abcam, Cambridge, MA). Immunostaining with the phospho-specific antibodies require great care as they proved extremely sensitive to fixation and pH. Over-fixation and improper pH monitoring will fail to stain.

### Confocal Microscopy and post-processing

Confocal: Microscopy was performed using an LSM SP8 confocal microscope (Leica) using either a 20× 0.7 NA multi-immersion objective or a 63× 1.3 NA glycerol immersion objective. The Mark and Find mode was used to automate the collection of images encompassing the entire limbus and generated a folder full of Z stacks at various individual overlapping positions along the limbus. For collections that were intended for colocalization, imaging was done after automatically optimizing the XYs format to set the optimal number of pixels to meet the Nyquist criterion for best lateral resolution for NA of objective and emission wavelength.

Deconvolution: Deconvolution was performed using Huygens Deconvolution software (Scientific Volume Imaging, Netherlands). Point spread function (PSF) was calculated automatically based on microscopic parameters and model of microscope. Classical maximum likelihood estimation method was used to perform deconvolution. 3D rendering: Individual confocal Z stacks (.lsm files) were processed directly using Imaris 9.3 (BitPlane AG, Zurich, Switzerland). Either the maximum intensity projection setting or the blend setting in Surpass mode of Imaris was used for 3D rendering. Images were oriented so that structures of interest were visible. Multiposition Z stacks generated using the Mark and Find setting on the confocal microscope were imported into Imaris and converted into Imaris files (.ims files). The resulting Imaris files were then stitched to generate a comprehensive Z stack encompassing the limbus using XuvTools ^81^. Deconvolution was performed using Huygens Deconvolution software using the Classic Maximum Likelihood Estimation algorithm. Colocalization of channels was performed using the colocalization function of Imaris on deconvolved images. Thresholding was performed to remove the background signal prior to performing colocalization. Thresholding varied slightly between different primary antibodies. The colocalization percentages were recorded and graphed, and statistics were obtained from using PlotsOfData ^82^. The colocalized channel was created in Imaris. Images were captured after rendering using the snapshot feature of Imaris and adjusted for levels (experimental group treated identically), and assembled in Adobe Photoshop CS6 (Adobe, San Jose, CA). Controls and experimental image sets were treated identically for all the above computational processes.

#### Electron Microscopy

Mice were euthanized and eyes immediately were enucleated and fixed with 0.8% paraformaldehyde and 1.2% glutaraldehyde in 0.1 M phosphate buffer pH 7.2 at 4°C. The eyes were fixed for 1 h and the anterior segment removed and cut into 1 × 2 mm blocks that included cornea, iris, TM and ciliary body. Fixation was continued at 4°C for 12 h and the tissues were washed in phosphate buffer and then post-fixed with 1% osmium tetroxide, dehydrated and embedded in Embed-812 resin. Sections of 1 µm were cut and stained with Toluidine Blue O for orientation and determining if open SC was present and morphology of the limbus was intact. Ultrathin sections of 80-100 nm were cut and collected on TEM grids and stained with uranyl acetate and lead citrate. Images were collected on a JOEL JEM-1230 electron microscope operating at an accelerating voltage of 80kV and captured on a NAOSPRT15 camera at an exposure time of 2000 msec.

#### Immunoblotting

Protein lysate was prepared from pools of 6 limbal strips from B6 and B6. *Fyn^-/-^* eyes (3 sets). Briefly, limbal strips were dissected from eyes on ice and placed in liquid nitrogen. The frozen limbal strips were crushed into powder using a pestle. The resulting powder was extracted with 200µl ice cold RIPA buffer (MILLIPORE SIGMA, Burlington, MA) with Halt™ Protease and Phosphatase Inhibitor Cocktail (1:100,78440, Thermo Fisher Sci, Waltham, MA) for 30 min at room temperature and then a further 30 min on ice. The lysates were sonicated briefly and spun down at 14,000Xg for 30 min. The supernatant was diluted 1:1 with Laemmli Sample Buffer (1610737, BIO-RAD, Hercules, CA) and heated to 80°C for 30 min.

30µl of the B6 and B6. *Fyn^-/-^* lysate was loaded (4 sets per gel) on 4-20% TGX precast Mini-PROTEAN gels (4561103, BIO-RAD, Hercules, CA) and run at 100 V. The gel was transferred to PVDF membrane using an iBLOT dry transfer system (Thermo Fisher Sci, Waltham, MA). The blots were cut into strips to give B6 and B6 and B6. *Fyn^-/-^* lysate paired lanes. The membrane was blocked using 5% bovine serum albumin in Tris-buffered saline plus 0.1% Tween 20 (TBST) for 1h at RT. The following antibodies were used for immunoblotting: Goat IgG against VE-cadherin (1:500, AF1002, R&D Systems, Minneapolis, MN), pY658 VEC (1:250, 44-1144G, Thermo Fisher Sci, Waltham, MA), FYN (1:500, HPA023887, SIGMA-ALDRICH, Burlington, MA), pY418SRC (1:500, ab4816, Abcam, Cambridge, MA) and GAPDH (1:500,). Appropriate horse radish peroxidase-conjugated antibodies were used at a dilution of 1:2500. The blots were developed using SuperSignal™ West Pico PLUS Chemiluminescent Substrate (34580, Thermo Fisher Sci, Waltham, MA). The blots were imaged on the Azure c600 Ultimate Western system. Images were further processed (levels) and assembled in Adobe Photoshop.

### IOP measurements

IOP was measured using the microneedle method as previously described in detail ^83, 84^. Briefly, mice were acclimatized to the procedure room and anesthetized via an intraperitoneal injection of a mixture of ketamine (99 mg/kg; Ketathesia, Henry Schein, Dublin, OH, USA) and xylazine (9 mg/kg; Anased, Akorn Inc, Gurnee, IL, USA) immediately prior to IOP assessment, a procedure that does not alter IOP in the experimental window ^84^.

### Pressure perfusion of mouse eyes

Pressure clamping or perfusion of eyes: Mice were anesthetized using isoflurane (5% for induction, 2% for maintenance) and kept warm using a 37°C heated mount. PBS was placed on the eye to prevent drying. The eye was cannulated using a beveled 60 µm pulled glass needle connected to a pressure sensor and a gravity-driven hydrostatic column filled with PBS. Following cannulation, the column was adjusted to induce the raised pressure (16mm Hg, 25 mmHg, or 45 mmHg) in the eye for 45 minutes. These pressures were picked based on an earlier work showing effects of pressure on effects on SC junctions ^20^.

To label the path of AQH outflow, 0.5 mg/ml of Alexa 594 conjugated GSII lectin (L21416, Thermo Fisher Sci, Waltham, MA) in PBS was perfused at the desired pressure.

Rapid fixation of eyes under pressure: After clamping eyes at the desired pressure for 45 min, the eyes were numbed with proparacaine for one minute under pressure with the needle still inserted in the eye. Then to fix the eyes under pressure, 10µl of a solution of 1.1% acrolein and 2% paraformaldehyde (PFA) in PBS was applied topically to the eye. After 5 min, the eye was washed with PBS. The mouse was euthanized, and the eye was enucleated and placed in 4% PFA for one hour at 4 °C. The eye was then subjected to several room-temperature washes. First were two five-minute washes in PBS, followed by a 20-minute wash in 1% sodium borohydride in PBS, and finally, six more five-minute washes in PBS before immunofluorescence.

#### In vivo AQH outflow measurement in mice

Outflow facility was measured by a two-step pressure method and calculated based on the Goldman equation F = (IOP-EVP) x C + Fu, where C= conventional outflow facility, F = total outflow rate, IOP = is intraocular pressure, EVP = Episcleral venous pressure, and Fu = unconventional outflow ^85^. The facility of the mouse eye was calculated by measuring flow rates of fluid in a specified time at two different pressures. Mice were anesthetized with isoflurane (5% for induction, 2% for maintenance) and secured on a heated mount (37°C) with a nose cone for isoflurane delivery. After placing a drop of PBS, on the cornea, an eye was cannulated using a beveled 60 µm glass needle mounted on a capillary holder attached to a pressure transducer (DTXPlus TNF-R; Argon Medical Systems, Plano, TX, USA; a range of -30 mmHg to 300 mmHg and hysteresis and sensitivity variations is no more than 2% of the reading or ±1 mmHg, whichever is greater, over the operating range). The pressure transducer, in turn, was connected by a stopcock and a short length of pressure-resistant tubing to a gravity-driven pressure (hydrostatic) column with an in-line flow sensor (SLG-0150; Sensirion AG, Switzerland)^86^. The pressure transducer is connected to a computer via an amplifier and analog-to-digital converter. The pressure measurements were made with custom-built software ^74^. The flow sensor is connected to the computer through a USB cable and software provided by Sensirion was used to measure flow.

The hydrostatic column, pressure and flow sensors, stopcock, and needle were all filled with PBS. The stopcock was kept closed prior to cannulation. It was opened following cannulation to induce a pressure of 20 mmHg in the eye. Following five minutes of equilibration, the total volume of liquid that flowed from the column over 10 minutes was measured. The pressure was then raised to 30 mmHg, and the five minutes of equilibration and 10 minutes of measurement were repeated. C, the facility was calculated by dividing the difference between two flow rates by the difference between the corresponding pressures and time elapsed to give a value in nl/min/mmHg. The AQH outflow data had a normal distribution based on Shapiro-Francia (p=0, W’=0.0511) and Anderson-Darling (p=0, W=0.3683). The data was graphed using R studio and ggplot2.

#### Statistical analysis

Statistical analysis of data, including t-test, One-way ANOVA, was done using Microsoft Excel and R. The median absolute deviation (MAD) is a robust measure of the variability of quantitative data and was used to judge the dysregulation of IOP and AQH outflow facility. Variance and quartiles were determined by Microsoft Excel formula. Some summary statistics were also obtained from PlotsOfData ^82^. Pearson’s correlation graphing, correlation coefficient and p values were obtained using R.

## SUPPLEMENTARY FIGURE LEGENDS

**Figure S1.**
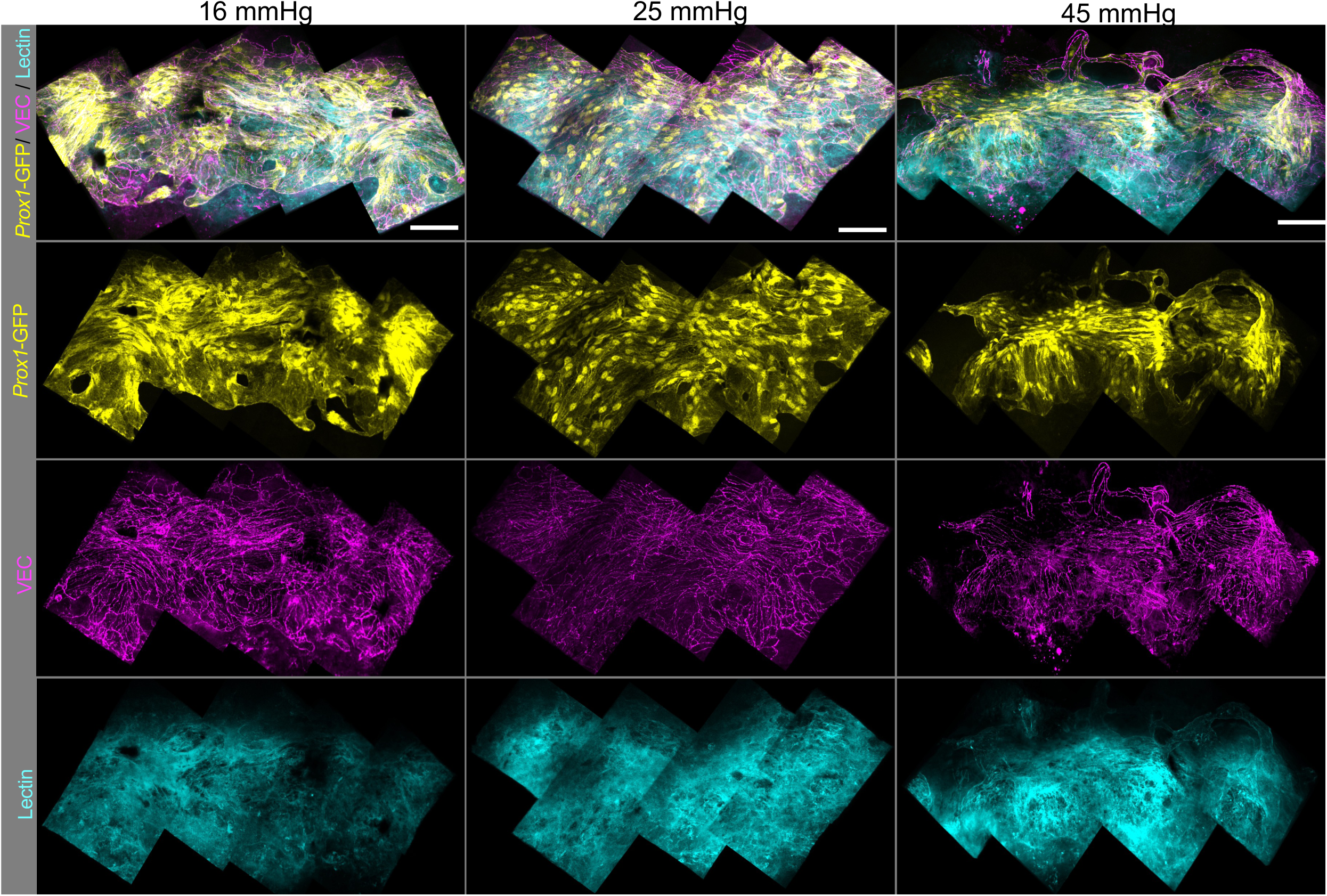
Lectin tracer levels increase at junctions marked by VE-CADHERIN in the SC inner wall with increasing IOP. *Prox1*-GFP eye perfused with lectin at 16,25 and 45 mmHg. This figure is an addendum to Figure 2B and shows the SC inner wall, VEC and Lectin channels plus the overlay. Scale, 25µm.

**Figure S2.**
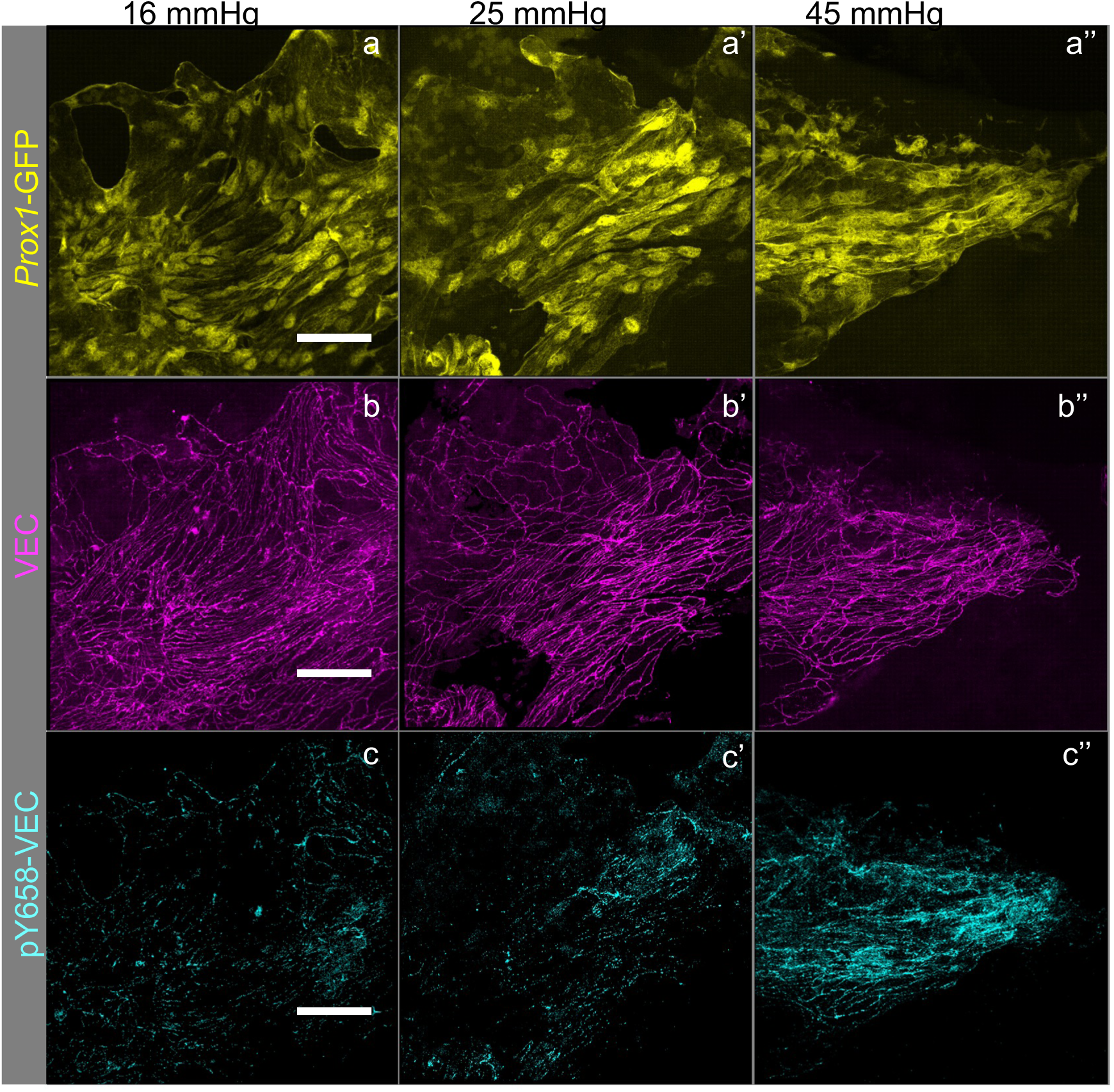
pY658 tyrosine phosphorylation in the VE-CADHERIN cytoplasmic domain increases with increasing IOP. *Prox1*-GFP eye pressure clamped at 16 mmHg, 25 mmHg, and 45 mmHg. **a-c**, the inner wall of SC showing GFP and VEC labeling, **a’-c’** VEC and **a’’-c’’** localization of pY658VEC at cell-cell junctions at the indicated pressure. Each dot represents data from a 246µm length of SC from each quadrant. N=6 eyes/ pressure. Scale, 20µm.

**Figure S3.**
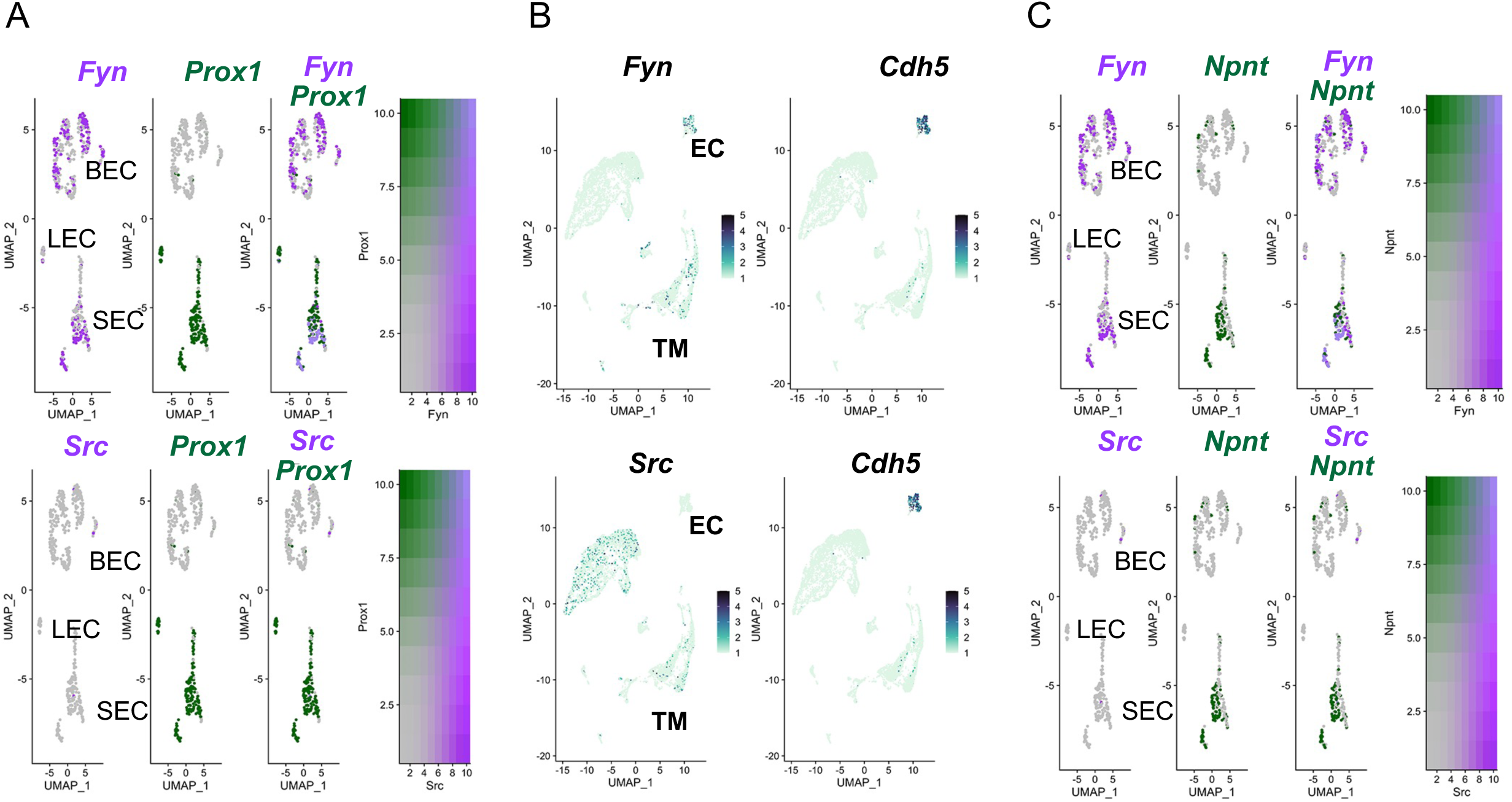
Feature plot representation of limbal endothelial cells analyzed by single cell RNA sequencing. A. *Fyn,* but not *Src, is* expressed in SECs (which also express *Prox1*) and BECs. B. *Fyn* has very minimal expression in TM cells, while SRC is more highly and commonly expressed in TM cells. C. *Fyn* but not *Src* localizes to SC inner wall cells that express *Npnt*. (Our single cell data show that *Npnt* is marker for IW cells ref). *Cdh5* is a marker for endothelial cells. The EC cluster in B includes all EC types and is subclustered into its distinct EC types in A and C.

**Figure S4.**
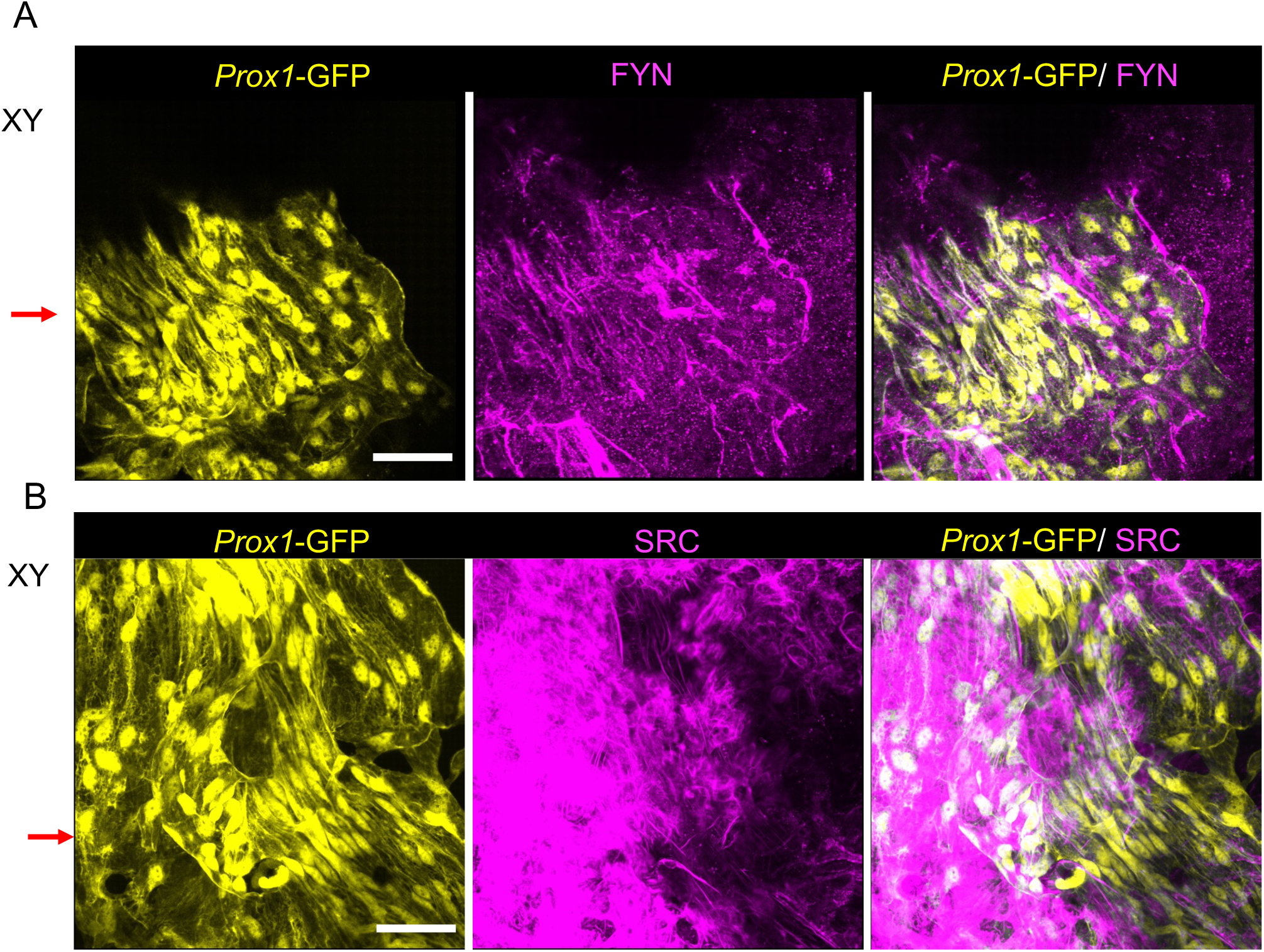
Localization of SFKs in the SC and the TM. Localization of FYN. FYN localization (magenta) in *Prox1*-GFP (yellow) eyes where XY image corresponds to XZ images in Fig 6. XZ optical sections were sliced along the section of XY. N=4 eyes. Scale, 20µm.

**Figure S5.**
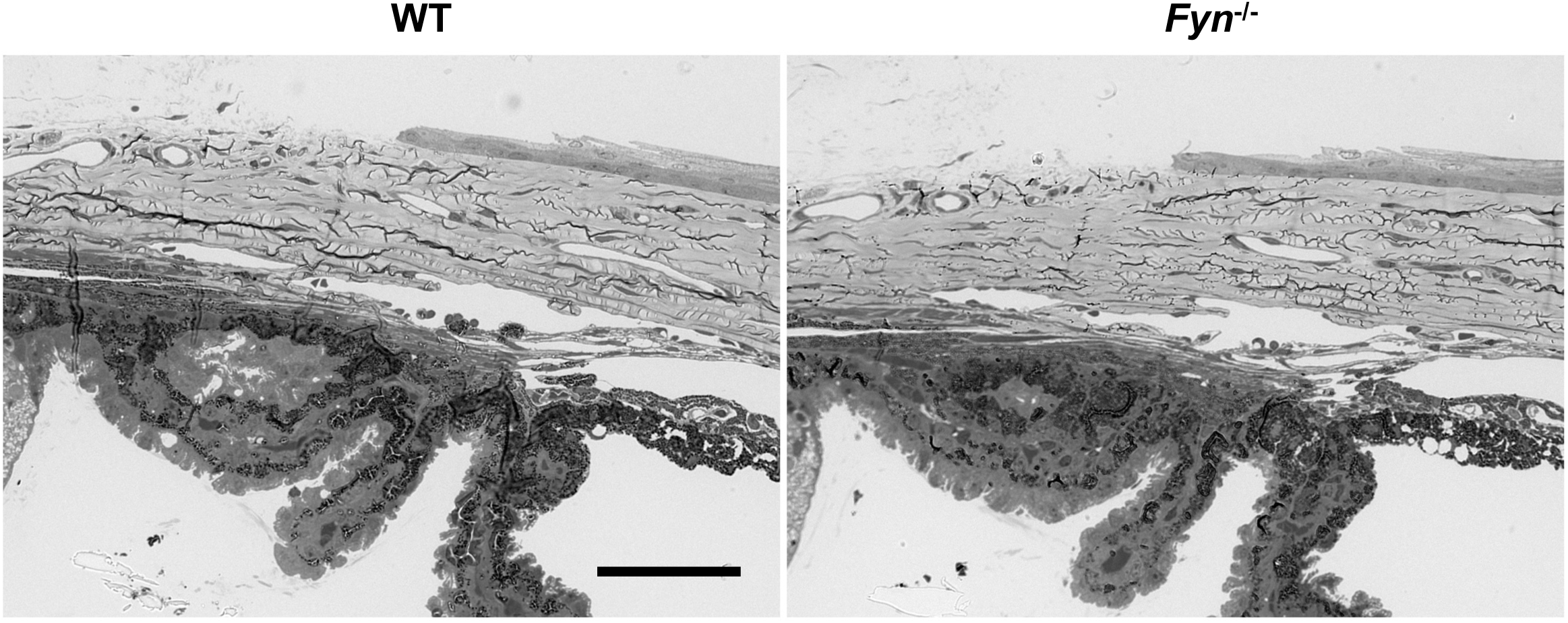
SC and TM show comparable morphology in wild type and FYN null eyes. Semithin sections of the iridocorneal region of the eyes from wild type and Fyn null eyes are shown. Scale 25µm.

**Figure S6.**
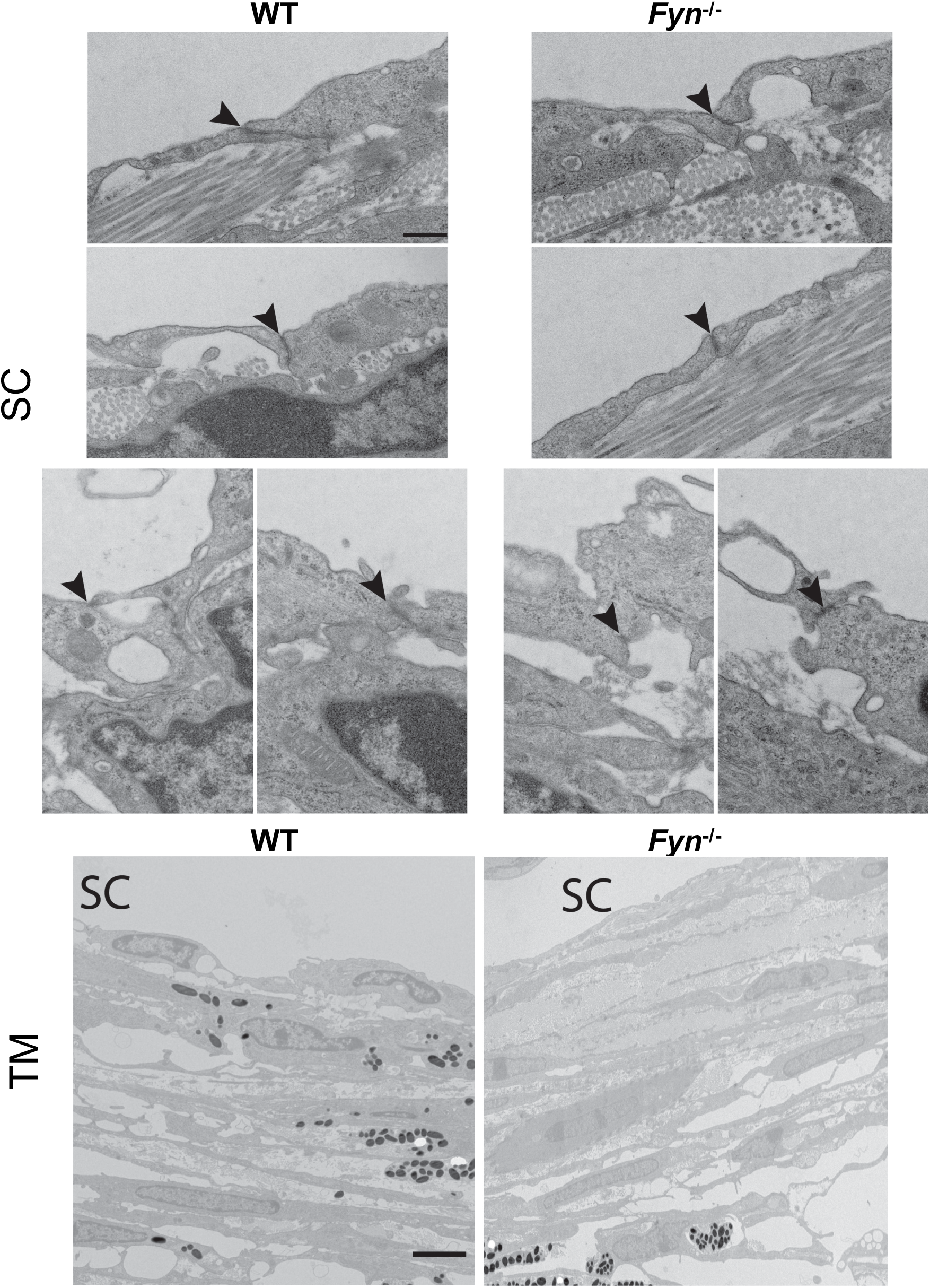
SC and TM show comparable morphology in wild type and FYN null eyes using transmission electron microscopy. Top panels show inner wall cells of the SC. Cell junctions are indicated by arrowheads. Bottom, TM. Scale, Top 400nm, bottom 4µm.

**Figure S7.**
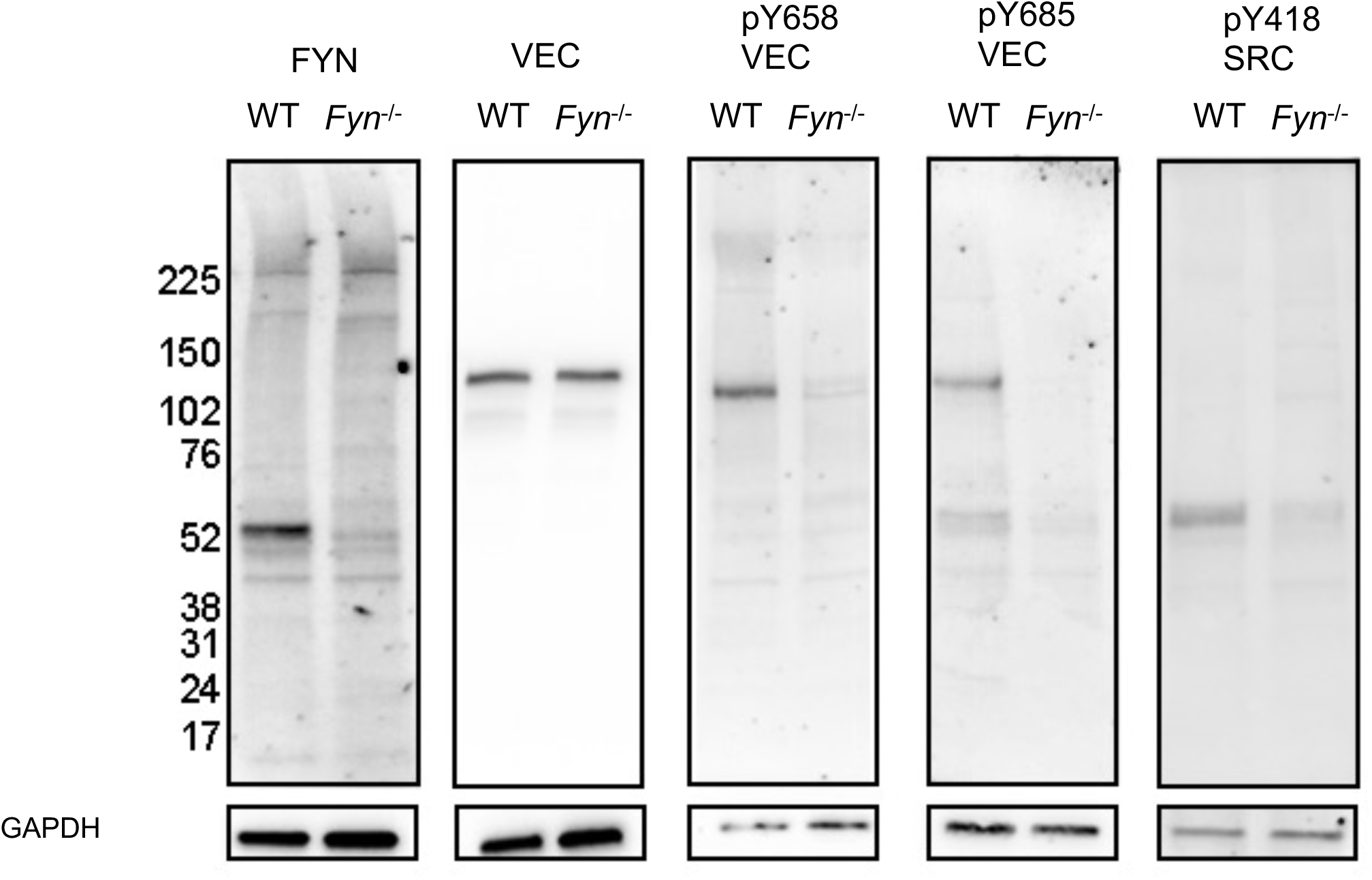
Loss of FYN results in the reduction of phosphorylation of VEC and SFK. Comparison of B6 and B6.*Fyn^-/-^* limbal strip lysates (pool of 6 eyes) by immunoblotting shows the reduction of pY658VEC and pY418 SRC levels with loss of FYN. Antibodies used for immunoblotting are shown on top of the blots. The first immunoblot shows VEC levels are not altered by loss of FYN in mouse limbus. Also shown are GAPDH blots for each of the immunoblots. Based on normalization with GAPDH, FYN protein levels were reduced by 90%, pY658VEC by 98%, pY685 by almost 100%, and pYSRC418 by 72%. VEC was unchanged.

**Figure S8.**
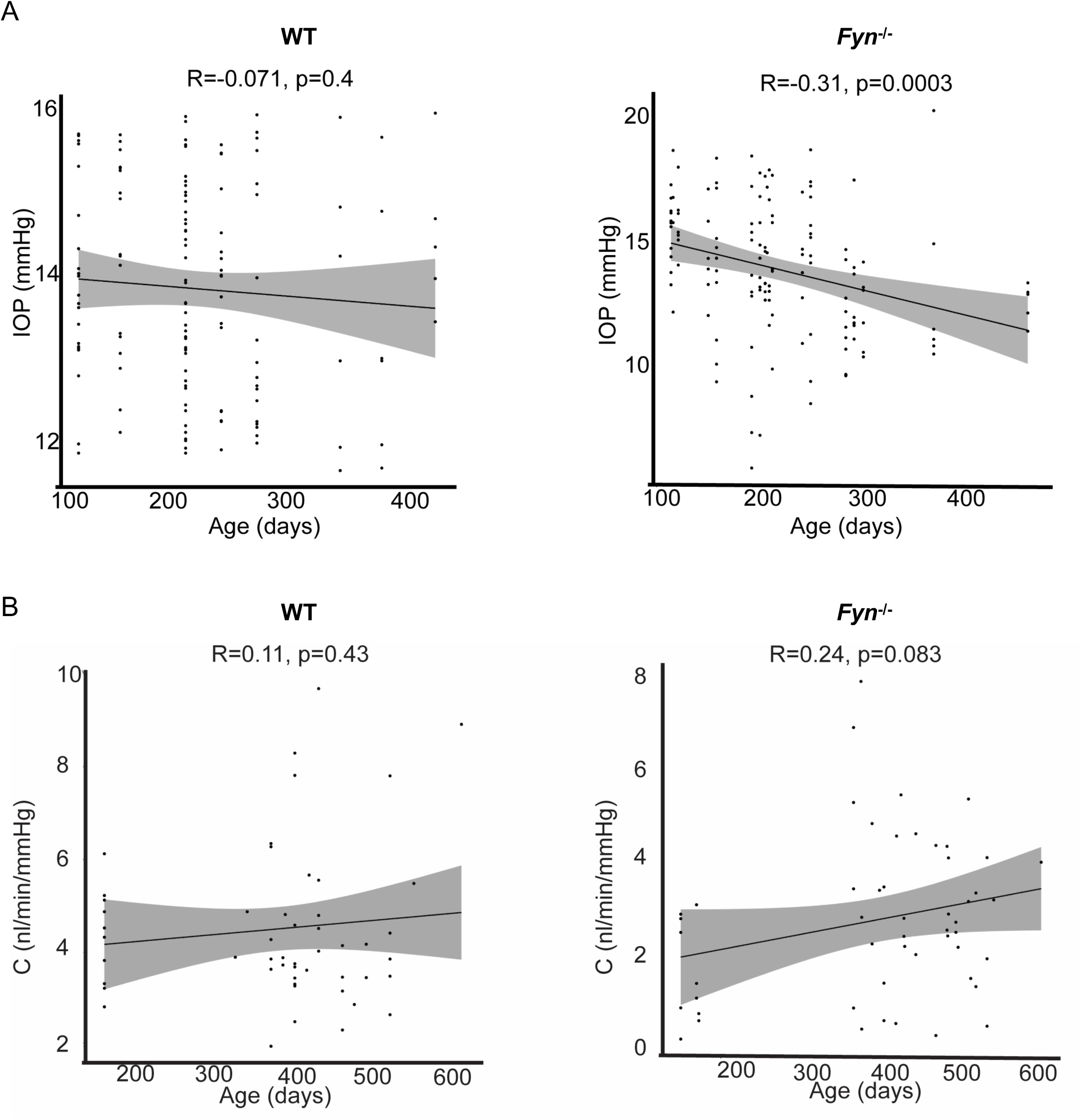
IOP and outflow show no correlation with age. Plots show results from a Pearson’s correlation analysis. **A**. IOP correlates poorly with age in both genotypes. **B.** Outflow facility C correlates poorly with age in both genotypes. Correlation coefficients and p values are shown above the plots.

